# Structural basis of RNA recognition by the SARS-CoV-2 nucleocapsid phosphoprotein

**DOI:** 10.1101/2020.04.02.022194

**Authors:** Dhurvas Chandrasekaran Dinesh, Dominika Chalupska, Jan Silhan, Vaclav Veverka, Evzen Boura

## Abstract

Severe acute respiratory syndrome coronavirus 2 (SARS-CoV-2) is the causative agent of the Coronavirus disease 2019 (COVID-19) which is currently negatively affecting the population and disrupting the global economy. SARS-CoV-2 belongs to the +RNA virus family that utilize single-stranded positive-sense RNA molecules as genomes. SARS-CoV-2, like other coronaviruses, has an unusually large genome for a +RNA virus that encodes four structural proteins – the matrix (M), small envelope (E), spike (S) and nucleocapsid phosphoprotein (N) - and sixteen nonstructural proteins (nsp1-16) that together ensure replication of the virus in the host cell. The nucleocapsid phosphoprotein N is essential for linking the viral genome to the viral membrane. Its N-terminal RNA binding domain (N-NTD) captures the RNA genome while the C-terminal domain anchors the ribonucleoprotein complex to the viral membrane via its interaction with the M protein. Here, we characterized the structure of the N-NTD and its interaction with RNA using NMR spectroscopy. We observed a positively charged canyon on the surface of the N-NTD lined with arginine residues suggesting a putative RNA binding site. Next, we performed an NMR titration experiment using an RNA duplex. The observed changes in positions of signals in the N-NTD NMR spectra allowed us to construct a model of the N-NTD in complex with RNA.

## Introduction

Positive-sense, single-stranded RNA viruses pose a global threat to human health as documented by the recent Zika virus outbreak and, more importantly, by the current COVID-19 pandemic caused by the severe acute respiratory syndrome coronavirus 2 (SARS-CoV-2) of the *Coronaviridae* family. SARS-CoV-2 earned its name due to the similarity of its genome to that of the SARS virus. Another deadly virus, the MERS (Middle East respiratory syndrome) virus is also a member of the *Coronaviridae* family, demonstrating how dangerous its members can be to humans. However, the recent COVID-19 pandemic is noteworthy. It is the most dangerous coronavirus to date, causing thousands of deaths, overwhelming the global health care system capacity, disrupting our everyday lives and causing enormous economic damage. It also reminds us of the vulnerability of our civilization to pathogens and highlights the importance of antiviral research.

Coronaviruses have the largest known genomes (up to 32 kb) among +RNA viruses, and they encode four structural and sixteen non-structural proteins. The non-structural proteins carry all of the enzymatic activities important for the viral replication, mostly associated with RNA replication. The genome SARS-CoV-2 also encodes an RNA-dependent RNA-polymerase complex (nsp7, nsp8 and nsp12), RNA capping machinery (nsp10, nsp13, nsp14 and 16) and additional enzymes such as proteases (the nsp3 PL^pro^ and the nsp5 3CL^pro^) which cleave viral polyproteins and/or impede innate immunity (1). The four structural proteins together with the viral +RNA genome and the envelope constitute the virion. The matrix (M), small envelope (E), and spike (S) proteins are embedded within the lipid envelope. The fourth structural protein, the nucleocapsid phosphoprotein (N), physically links the envelope to the +RNA genome. It consists of an N-terminal (NTD) and a C-terminal (CTD) domain (Figure 1). Both domains are capable of RNA binding. In addition, the CTD serves as a dimerization domain and binds the matrix protein forming the physical link between the+ RNA genome and the envelope. The SARS N protein has also been shown to modulate the host intracellular machinery and plays regulatory roles during the viral life cycle (2). In light of the genomic similarities between SARS-CoV and SARS-CoV-2, it is reasonable to expect the N protein to function in a similar way.

**Figure 1.**
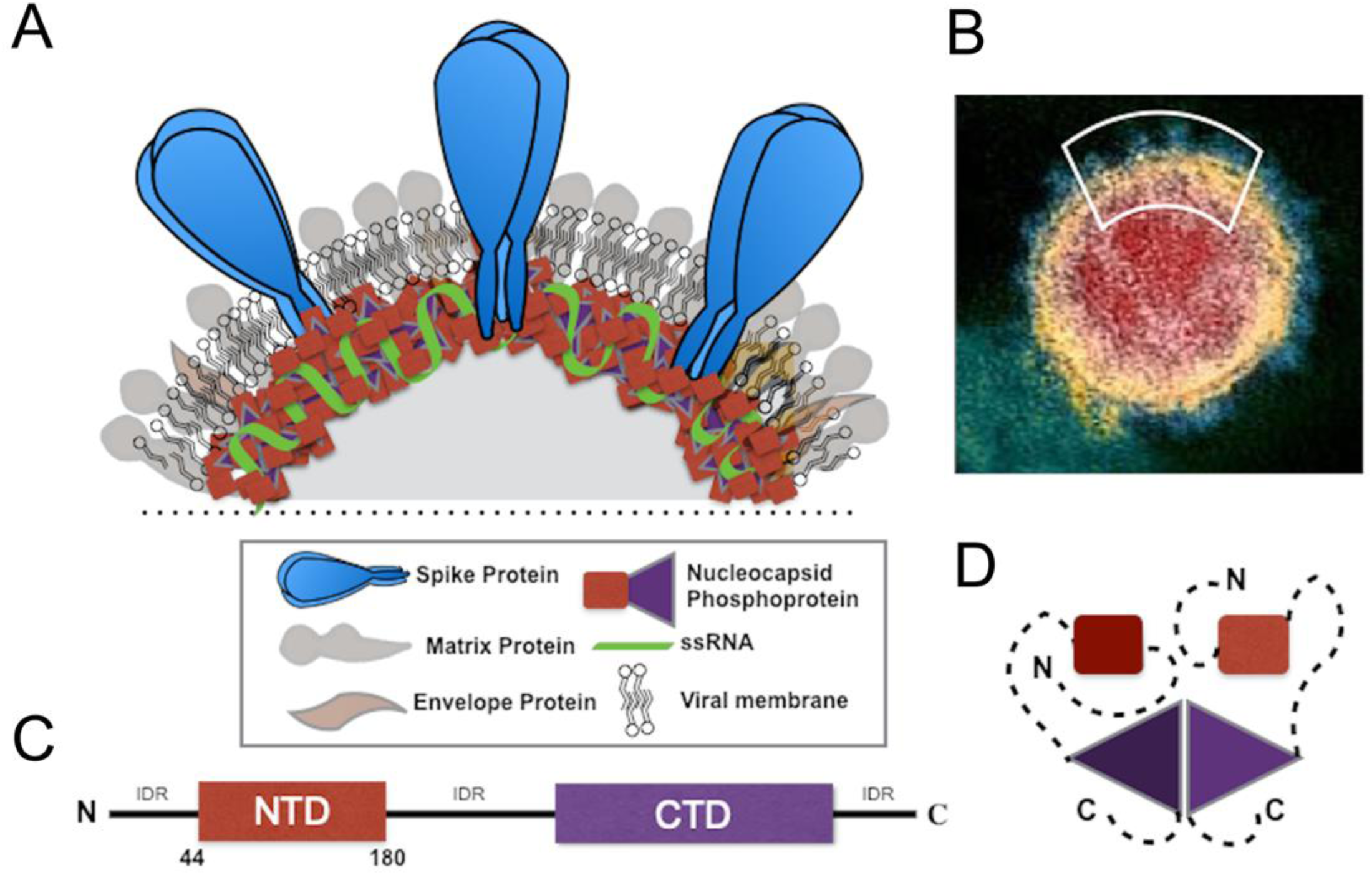
SARS-CoV-2 virion and model of structural proteins. (A) Enlarged view of the viral membrane showing the four structural proteins: spike – S, membrane – M, envelope – E, and nucleocapsid phosphoprotein – N, along with viral membrane and the RNA genome. (B) Transmission electron microscopic image of a single SARS-CoV-2 viral particle (NIH, NIAID-RML). (C) Domain organization of the full length N-protein. (D) Schematic organization of the N protein dimer formed through the CTD domain (the N-NTD is shown in red and the N-CTD in lavender)

All the SARS-CoV-2 proteins are potential drug targets (3) and a detailed understanding of their functions is therefore of utmost importance. An experimental treatment using remdesivir, which has been tested in several countries strongly affected by COVID-19, targets the RNA-dependent RNA-polymerase nsp12, as does favipiravir (also known as T-705), which has been approved for clinical use in Japan and China. Both remdesivir and favipiravir are nucleotide analogs. However, unlike most RNA viruses, SARS-CoV-2 encodes an exonuclease (a second enzymatic activity of the RNA capping factor nsp14) presumably capable of repairing mismatches in newly synthesized double-stranded RNA. Additional antiviral compounds might be necessary to simultaneously target several viral proteins and create a trap that the virus cannot escape by mutation. In any case, drugs targeting proteins other than the RNA polymerase are urgently needed. In this study, we have analyzed in detail the structure of the N protein NTD (N-NTD) and its interaction with RNA using protein NMR. Based on this data we generated a model of the N-NTD and its complex with RNA that illustrates how the N protein recognizes RNA and provides a basis for rational drug design.

## Results

Here, we report the NMR structure of the SARS-CoV-2 nucleocapsid phosphoprotein N-terminal RNA binding domain N-NTD. The structure revealed an overall right hand-like fold composed of a β-sheet core with an extended central loop. The core region adopts a five-stranded U-shaped right-handed antiparallel β-sheet platform with the topology β4-β2-β3-β1-β5 flanked by two short α-helices (α1 occurs before the β2 strand and α2 after the β4 strand). A prominent feature of the structure is a large protruding loop between β2-β3 that forms a long basic β-hairpin (β2’ and β3’) (Figure 2). This long β-hairpin is reminiscent of a finger and composed mostly of basic amino acids residues, and we therefore refer to it as a basic finger (Figure 2). This basic finger extends from the β-core structure that we refer to as a palm. Analysis of the electrostatic potential of the N-NTD revealed a highly positively charged cleft between the basic finger and the palm creating a putative RNA binding site in the hinge/junction region between the palm and basic finger. Our NMR analysis is consistent with recent X-ray analysis (PDB IDs: 6M3M and 6VYO). In addition, the NMR structure revealed that the basic finger is highly flexible (Figure 2A) whereas in the crystal structures it is locked in one place by crystal lattice contacts.

**Figure 2.**
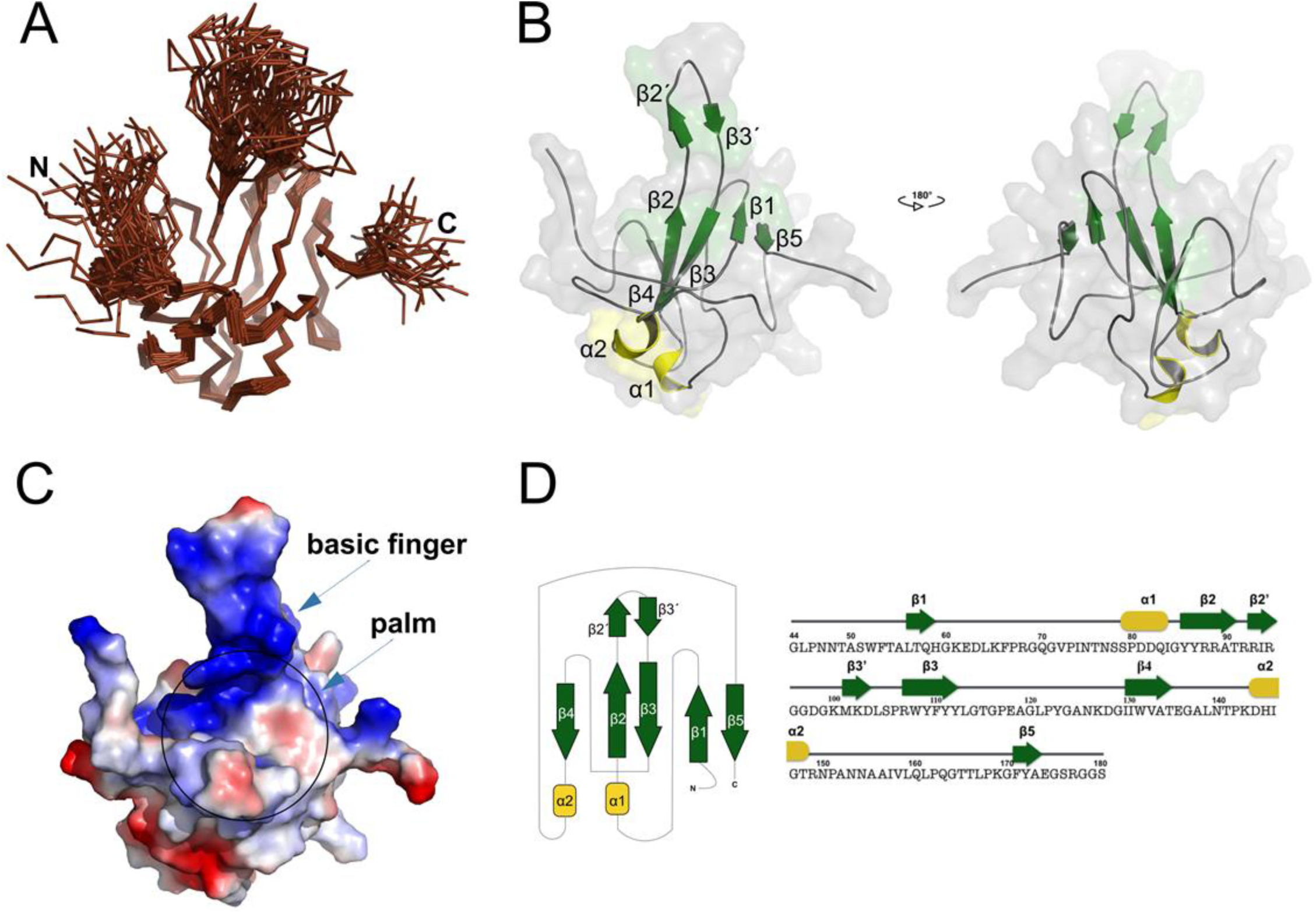
Solution structure of the SARS-CoV-2 N-NTD RNA binding domain. (A) Backbone representation of the 40 converged structures of N-NTD obtained by NMR spectroscopy. (B) Cartoon representation of the lowest energy structure (structural elements are highlighted in color: α1-α2 helices (yellow), *β*1-(*β2’-β3’*)-*β5* (green), and loops (gray)) show the overall U-shaped antiparallel β-sheet platform (the palm) and a protruding β-hairpin (the basic finger). (C) The N-NTD molecular surface electrostatic potentials revealed a basic patch extended between the finger and the palm, with positively charged surface shown in blue and negatively charged surface in red. (D) Topology diagram of the N-NTD.

The structure of the N-NTD suggests a putative RNA binding site. Intrigued, we aimed to obtain an experimental model of the N-NTD in a complex with RNA. We performed an NMR-based titration experiment using a short double-stranded RNA (5’-CACUGAC-3’ and 5’-GUCAGUG-3’). Basically, we were adding isotopically unlabeled RNA to the ^15^N/^13^C labeled protein and we followed changes in positions of assigned signals in NMR spectra (Figure 3) to reveal the molecular interface of the N-NTD:RNA complex. Overlay of the 2D ^15^N/^1^H HSQC and 2D ^13^C/^1^H HSQC spectra of a free and RNA bound N-NTD revealed residues that were significantly perturbed by RNA binding. All these significantly affected residues (A50, T57, H59, R92, I94, S105, R107, R149, Y172) are located in the basic finger or close to the junction between the basic finger and the palm as expected based on the analysis of the electrostatic potential.

**Figure 3.**
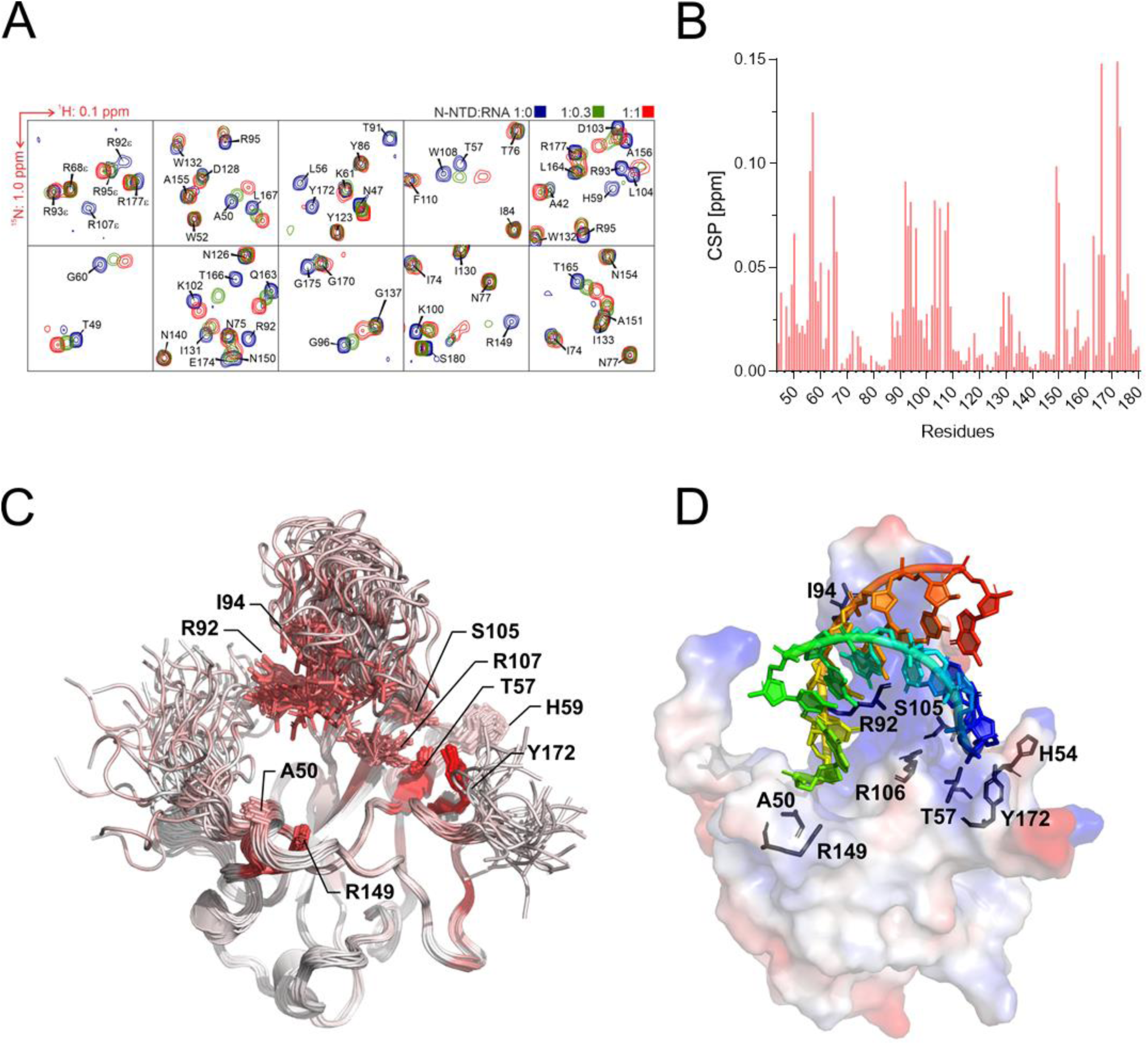
NMR-based mapping and a model of the N-NTD:RNA complex. (A) Representative regions from the 2D ^15^N/^1^H HSQC titration spectra illustrating effects of addition of the 7mer RNA duplex on backbone and side-chain N-NTD amide signals (arginine side-chains are labeled with ε – top left panel). The 50 µM ^15^N-labeled N-NTD protein construct was titrated with an increasing concentration of RNA. (B) Chemical shift perturbations of N-NTD residues upon binding of an equimolar concentration of 7mer RNA duplex. (C) The CSPs highlighted on as a red-color gradient on the ribbon representation of 40 converged structures of N-NTD. The majority of the perturbed residues form a large continuous patch from R92 to Y172 on one side of the positively charged cleft and a smaller patch formed by A50 and R149 on the opposite side of the cleft. The solvent exposed sidechains of the most perturbed residues are shown as sticks. (D) N-NTD:RNA complex. The RNA duplex is shown as a cartoon structure over the electrostatic surface of SARS-CoV-2 N-NTD and the interacting residues are highlighted as sticks.

We next used the experimental data to build an atomic model of the protein:RNA complex. We used the HADDOCK protocol for NMR-restraint driven docking simulations (4). As a starting conformation of the RNA we chose a short double helix based on the published crystal structure of a short RNA duplex (5). A detailed analysis of the chemical shift perturbations (CSP) mapped onto the solution structure of N-NTD provided a set of ‘active’ solvent-accessible residues on N-NTD that were expanded to surrounding ‘passive’ residues. The selection criteria for active residues were that their CSP values were more than one standard deviation from the average value (0.05 ppm) calculated for the entire set of CSPs and that their accessibility to solvent was more than 20% (6). Restraints for the RNA molecule were kept ambiguous to avoid potential bias. A standard docking protocol yielded a set of water-refined conformations for the protein:RNA complex that were clustered into several distinct classes. As expected, the RNA duplex molecule was bound in the positively charged cleft in all clusters. The most populated cluster also contained the fewest violations of experimental restraints and was therefore selected as a representative conformation for the N-NTD: RNA complex.

The structure revealed an RNA duplex bound to the positively charged canyon located between the basic finger and the palm of the N-NTD. A notable feature of the binding interface is its electrostatic potential. It is highly positive with several arginine residues (R92, R107, R149) that directly contact the RNA. Our model also predicts several hydrophobic interactions, such as one involving the side-chains of residues I94 and L104, which also contribute to RNA binding. Taken together our data provide a molecular snapshot of RNA recognition by the coronaviral N protein and suggest a hot spot located at the interface of the basic finger and the palm as a suitable target for targeting by small molecules.

## Discussion

Effective drugs are urgently needed to combat the COVID-19 disease. Most patients are not given any drug and the treatment relies on curing the symptoms. The most promising drug is remdesivir, a nucleotide analog that targets the viral RNA-dependent RNA-polymerase (RdRp). Viral polymerases are certainly good targets for antiviral compounds as these enzymes are absolutely vital for any +RNA virus. However, every viral protein is a potential target for antiviral compounds and an effective treatment may require several active compounds, each targeting a different protein at the same time. This approach, known as HAART (highly active antiretroviral therapy), has proven effective in the case of HIV, which is another virus with an RNA genome. In this study, we structurally characterized the N protein, a structural protein that is essential for assembly of the virion. We obtained an atomic model of the N protein N-terminal domain in an unliganded form and in complex with an RNA duplex. The structure revealed a right hand fold featuring a prominent basic finger protruding from the palm. Analysis of its electrostatic potential (Figure 2C) revealed highly positively charged canyon that is situated in the interface between the basic finger and the palm subdomain and constitutes a putative RNA binding site. We performed an NMR titration experiment to obtain experimental proof of the RNA binding site. An overlay of the ^15^N/^13^C labeled protein NMR spectrum in the absence of ligand and in complex with RNA revealed amino acid residues with large chemical shifts upon the addition of RNA (Figure 3). Not surprisingly, all these residues are located in or in close proximity to the basic canyon, confirming the canyon as the RNA binding site. To illustrate how the coronaviral N-NTD recognizes RNA we built an atomic model of the N-NTD:RNA complex using the NMR titration data as a restraint for computer simulations in HADDOCK. The model reveals a hotspot on the surface of the N-NTD between the finger and palm subdomains that is essential for RNA binding and suggests this site as an ideal place for targeting by small molecules.

## Materials and Methods

### Protein expression, and purification

N-terminal domain of the SARS CoV-2 N protein (N-NTD, residues 44-180) was expressed as a fusion protein with 6xHis tag followed by cleavage site for TEV protease on its N-terminus. *Escherichia coli* BL21(DE3) expressing the protein were grown in minimal media containing ^15^NH_4_Cl and [U-^13^C]glucose. The culture was lysed by sonication in lysis buffer (50 mM Tris pH 8.0, 500 mM NaCl, 20 mM imidazole, 10% glycerol, 3mM β-mercaptoethanol) and the lysate was cleared by centrifugation. His-tagged protein was purified from the supernatant by affinity chromatography on a Nickel-NTA column (Machery-Nagel) according to manufactur’s instruction, the 6x His tag was cut off by the TEV protease and the protein was further purified by size-exclusion chromatography on a Superdex 75 HiLoad 26/60 column (GE Healthcare) in SEC buffer (20 mM Na_2_HPO_4_, 50 mM NaCl, 0.01% NaN_3_, pH 5.5). Purity of the protein was checked using SDS-PAGE. Protein was concentrated to 1.19 mM and used for NMR experiments. For further NMR measurements of binding RNA, the N-NTD was diluted to 300 μM and flash frozen in liquid nitrogen and stored in -80°C until needed.

To examine the RNA binding mode of the N-NTD we used a commercially available 7mer RNA duplex that was prepared by annealing of RNA oligonucleotides 5’-CACUGAC-3’ and 5’-GUCAGUG-3’ (Sigma). The RNA oligonucleotides were mixed in a molar ratio 1:1 at the final concetration 200 μM of each oligonucleotide and water supplemented with 50 mM NaCl. The mixture was incubated at 60°C for 15 min and then cooled slowly at 26°C. For RNA titration, annealed RNA was added to 50 μM protein in molar ratios 1:0.3125, 1:0.625, 1:1 and 1:2.

### NMR spectroscopy

NMR spectra were acquired at 25 °C on an 850 MHz Bruker Avance spectrometer, equipped with a triple-resonance (^15^N/^13^C/^1^H) cryoprobe. The sample volume was either 0.16 or 0.35 mL, in SEC buffer, 5% D_2_O/90-95% H_2_O. A series of double- and triple-resonance spectra (7, 8) were recorded to obtain sequence-specific resonance assignment. We used I-PINE assignment tool (9) implemented in NMRFAM-SPARKY (10) for initial automatic assignment. ^1^H-^1^H distance restraints were derived from 3D ^15^N/^1^H NOESY-HSQC and ^13^C/^1^H NOESY-HMQC, which were acquired using a NOE mixing time of 100 ms.

The structural calculation was carried out in Cyana (11) using NOESY data in combination with backbone torsion angle restraints, generated from assigned chemical shifts using the program TALOS+ (12). First, the combined automated NOE assignment and structure determination protocol (CANDID) was used for automatic NOE cross-peak assignment. Subsequently, five cycles of simulated annealing combined with redundant dihedral angle restraints were used to calculate a set of converged structures with no significant restraint violations (distance and van der Waals violations <0.5Å and dihedral angle constraint violations <5°). The 40 structures with the least restraint violations were further refined in explicit solvent using the YASARA software with the YASARA forcefield(13) and subjected to further analysis using the Protein Structure Validation Software suite (www.nesg.org). The statistics for the resulting structure is summarized in Table 1. The structures, NMR restraints and resonance assignments were deposited in the Protein Data Bank (PDB, accession code: 6YI3) and BMRB (accession code: 34511).

**Table 1.**
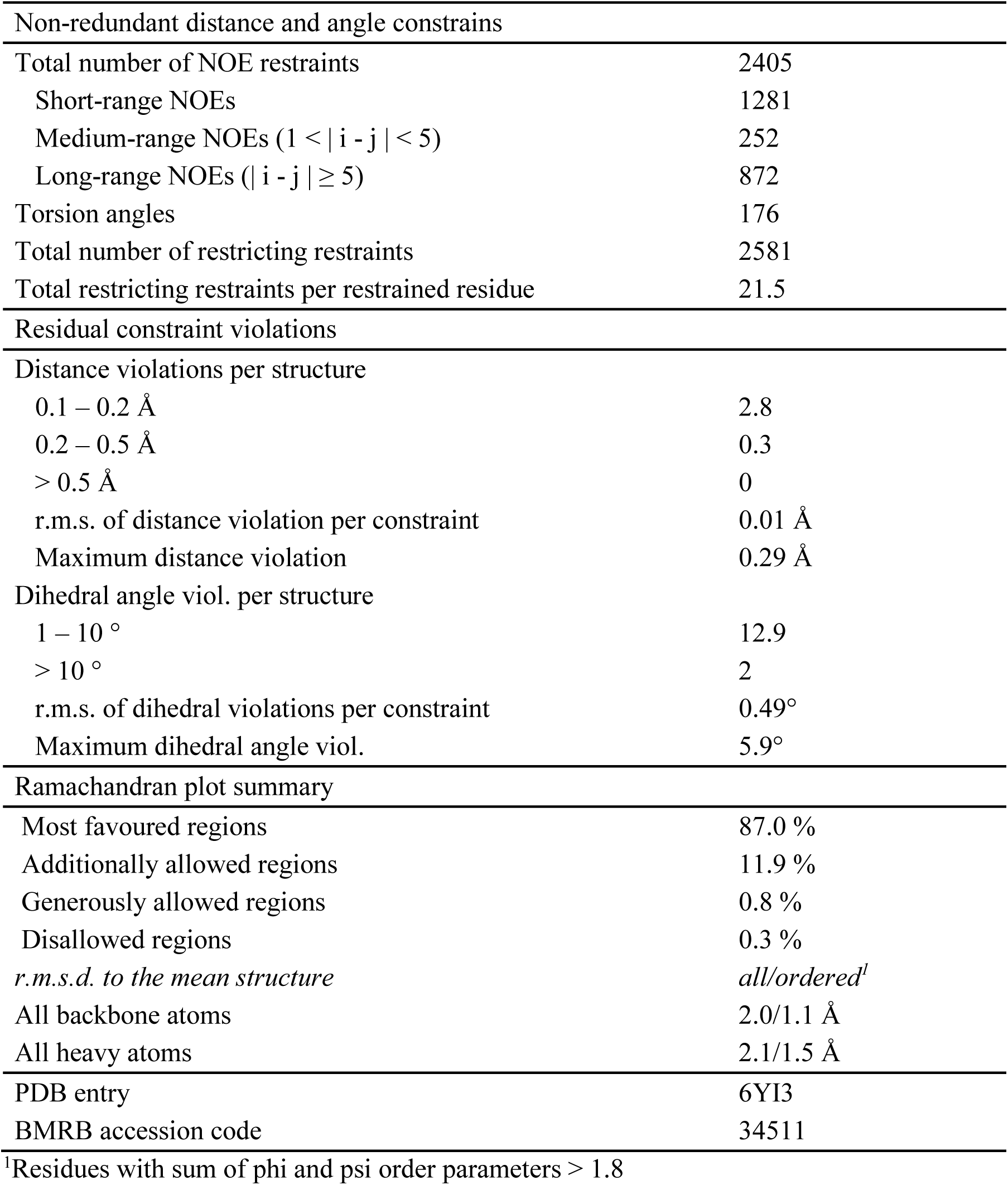
NMR Constraints and Statistics for the final set of structures.

To follow changes in the chemical shifts of a protein upon RNA binding, we calculated chemical shift perturbations (CSPs). The CSP of each assigned resonance in the 2D ^15^N/^1^H HSQC spectra of the protein in the free state is calculated as the geometrical distance in ppm to the peak in the 2D ^15^N/^1^H HSQC spectra acquired under different conditions using the formula: 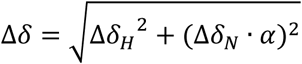, where α is a weighing factor of 0.2 used to account for differences in the proton and nitrogen spectral widths.(14)

### Molecular docking

The structure for the N-NTD in complex with the 7mer RNA duplex was calculated using HADDOCK (4). The RNA homology model was prepared by mutating the native 7mer RNA duplex (PDB 4U37) (5) in Pymol (The PyMOL Molecular Graphics System, Version 2.0 Schrödinger, LLC.) that was subsequently subjected to an energy minimization in YASARA (13). For the actual docking, we used a representative structure from the set of obtained structures and followed a standard protocol. As active were selected those N-NTD residues with CSP > 0.05 ppm and at least 20% solvent accessibility (A50, T57, H59, R92, I94, S105, R107, R149, A152 and Y172), while as passive were additionally selected adjacent solvent exposed residues (T49, T54, L55, R88, A90, K102, L104, Y109, Y111, P151, A155, A156, E174 and G175). On RNA side, all 14 nucleobases were defined as active for the experimentally driven docking protocol. In addition, in total three regions within the N-NTD were defined as fully flexible segments for the advanced stages of the docking calculation (the N-terminal G1-T9, the central loop I54-M61 and the C-terminal S136-S140). The final set of 200 water-refined structures was clustered using a Fraction of Common Contacts approach (15) with a default cut-off 0.75 and a minimal cluster size = 4. The resulting structures were sorted into 7 clusters and the most populated cluster (n = 30) that also exhibited the lowest interaction energy was selected for detailed analysis.

## Author Contributions

DCD, DC, JS and VV performed experiments, VV interpreted the NMR data and built the final model. VV and EB designed and supervised the project. VV and EB wrote the manuscript, all authors commented on the manuscript.

## Acknowledgement

The work was supported from European Regional Development Fund; OP RDE; Project: “Chemical biology for drugging undruggable targets (ChemBioDrug)” (No. CZ.02.1.01/0.0/0.0/16_019/0000729). The Academy of Sciences of the Czech Republic (RVO: 61388963) is also acknowledged. We are grateful to Edward Curtis for critical reading of the manuscript.

## Accession codes

The NMR restraints, resonance assignments and the structure were deposited in the PDB under accession code 6YI3 and in the BMRB database under accession code 34511).

## References

1. Snijder EJ, Decroly E, & Ziebuhr J (2016) The Nonstructural Proteins Directing Coronavirus RNA Synthesis and Processing. Advances in virus research 96:59–126.

2. Chang CK, Hou MH, Chang CF, Hsiao CD, & Huang TH (2014) The SARS coronavirus nucleocapsid protein--forms and functions. Antiviral research 103:39–50.

3. Gordon DE, et al. (2020) A SARS-CoV-2-Human Protein-Protein Interaction Map Reveals Drug Targets and Potential Drug-Repurposing. BioRxiv.

4. Dominguez C, Boelens R, & Bonvin AM (2003) HADDOCK: a protein-protein docking approach based on biochemical or biophysical information. Journal of the American Chemical Society 125(7):1731–1737.

5. Sheng J, Larsen A, Heuberger BD, Blain JC, & Szostak JW (2014) Crystal structure studies of RNA duplexes containing s(2)U:A and s(2)U:U base pairs. Journal of the American Chemical Society 136(39):13916–13924.

6. Wilkinson IC, et al. (2009) High resolution NMR-based model for the structure of a scFv-IL-1beta complex: potential for NMR as a key tool in therapeutic antibody design and development. The Journal of biological chemistry 284(46):31928–31935.

7. Renshaw PS, et al. (2004) Sequence-specific assignment and secondary structure determination of the 195-residue complex formed by the Mycobacterium tuberculosis proteins CFP-10 and ESAT-6. Journal of biomolecular NMR 30(2):225–226.

8. Veverka V, et al. (2006) NMR assignment of the mTOR domain responsible for rapamycin binding. Journal of biomolecular NMR 36 Suppl 1:3.

9. Lee W, et al. (2019) I-PINE web server: an integrative probabilistic NMR assignment system for proteins. Journal of biomolecular NMR 73(5):213–222.

10. Lee W, Tonelli M, & Markley JL (2015) NMRFAM-SPARKY: enhanced software for biomolecular NMR spectroscopy. Bioinformatics 31(8):1325–1327.

11. Herrmann T, Guntert P, & Wuthrich K (2002) Protein NMR structure determination with automated NOE assignment using the new software CANDID and the torsion angle dynamics algorithm DYANA. Journal of molecular biology 319(1):209–227.

12. Shen Y, Delaglio F, Cornilescu G, & Bax A (2009) TALOS+: a hybrid method for predicting protein backbone torsion angles from NMR chemical shifts. Journal of biomolecular NMR 44(4):213–223.

13. Harjes E, et al. (2006) GTP-Ras disrupts the intramolecular complex of C1 and RA domains of Nore1. Structure 14(5):881–888.

14. Veverka V, et al. (2008) Structural characterization of the interaction of mTOR with phosphatidic acid and a novel class of inhibitor: compelling evidence for a central role of the FRB domain in small molecule-mediated regulation of mTOR. Oncogene 27(5):585–595.

15. Rodrigues JP, et al. (2012) Clustering biomolecular complexes by residue contacts similarity. Proteins 80(7):1810–1817.

